# Predicting gene expression in the human malaria parasite *Plasmodium falciparum*

**DOI:** 10.1101/431049

**Authors:** David F. Read, Yang Y. Lu, Kate Cook, Karine Le Roch, William Stafford Noble

**Affiliations:** Department of Genome Sciences, University of Washington; Department of Cell Biology and Neuroscience University of California, Riverside; Department of Genome Sciences, Department of Computer Science and Engineering, University of Washington

## Abstract

Empirical evidence suggests that the malaria parasite *Plasmodium falciparum* employs a broad range of mechanisms to regulate gene transcription throughout the organism’s complex life cycle. To better understand this regulatory machinery, we assembled a rich collection of genomic and epigenomic data sets, including information about transcription factor (TF) binding motifs, patterns of covalent histone modifications, nucleosome occupancy, GC content, and global 3D genome architecture. We used these data to train machine learning models to discriminate between high-expression and low-expression genes, focusing on three distinct stages of the red blood cell phase of the *Plasmodium* life cycle. Our results highlight the importance of histone modifications and 3D chromatin architecture and suggest a relatively small role for TF binding in *Plasmodium* transcriptional regulation.

## 1 Introduction

*Plasmodium falciparum* is the deadliest species of malaria parasite, responsible for 445,000 deaths in 2016 [42]. As resistance to antimalarial drugs spreads, demand for novel antimalarials increases. Designing such novel drugs would require an improved understanding of the biology of this parasite. Currently, one of the primary open questions in *Plasmodium* biology is how the parasite maintains precise control of gene expression. The current work aims to address this question by constructing an accurate predictive model of *P. falciparum* transcription. The model accounts for the rich landscape of transcriptional control mechanisms in *Plasmodium* by incorporating five types of features, representing transcription factor (TF) binding, covalent histone modifications, nucleosome positioning, GC content, and chromatin 3D structure.

In many eukaryotes, TF binding within and around gene promoters is considered the dominant mechanism of gene expression control. However, in *Plasmodium*, several lines of evidence suggest that TF binding may be less central to transcriptional control. First, although major components of the general transcription machinery are present in the *Plasmodium* genome [17], a relatively small set of specific *Plasmodium* TFs (∼ 27) have been identified and validated in the parasite genome [17]. In comparison, the similarly sized genome of the yeast *S. cerevisiae* contains ∼ 170 specific TFs [18]. Second, among the subset of TFs whose binding affinities have been characterized via *in vitro* protein binding microarrays [19], only a handful display stage-restricted expression and play clear roles in regulating life cycle transitions. An example is PfAP2-G, which drives expression of gametocyte-specific genes [29, 52]. Third, a large number of *Plasmodium* genes are predicted by homology to function in the regulation of chromatin structure, mRNA decay, and translation [17], suggesting the importance of epigenetic and post-transcriptional regulation.

Among mechanisms for epigenetic regulation, patterns of covalent histone modifications are perhaps the most widely studied and understood. In this respect, some aspects of *P. falciparum* gene regulation are shared with other eukaryotes, including the presence of the typically heterochromatin-associated H3K9me3 histone modification at repressed *var* genes (referred to as virulence genes, for their role in parasite pathogenicity) [15] and depletion of promoter nucleosomes correlating with gene transcription [9]. On the other hand, *Plasmodium* epigenomic dynamics also exhibit notable deviations from those in commonly studied eukaryotes, such as abundant and broad distributions of activating histone marks [3, 35], an absence of H3K27me3 repressive marks [3], and genome-wide changes in nucleosome occupancy during the asexual cycle [9, 46]. These observations suggest that the parasite may make use of a “histone code” like other well-characterized eukaryotes, though the specific role of individual elements may differ.

In addition, empirical evidence suggests that gene regulation in *Plasmodium* occurs through changes in chromatin structure, including shifts in nucleosome occupancy at the local level and 3D positioning at larger scales. Nucleosome occupancy, as measured by MNase-assisted isolation of nucleolar elements (MAINE) and formaldehyde-assisted isolation of regulatory elements (FAIRE), or assay for transposase accessible chromatin with high-throughput sequencing (ATAC-seq), exhibit cyclic patterns that closely track changes in gene expression during the red blood cell (erythryocytic) stages of the parasite life cycle [46, 54]. In addition, 3D models of *Plasmodium* DNA based on Hi-C assays at multiple time points during the red blood cell [3] and transmission [8] stages of the parasite point to a “gradient” of expression across the nucleus, from a repressive center near the telomeres to an expressive center at the centromeres.

Based on the above evidence, we hypothesized that the cascade of transcripts observed throughout the red blood cell (erythrocytic) cycle is the result of a combination of transcription factor binding, histone modifications, and changes in chromatin structure. We further predicted than an integrated analysis of the relationships among transcription and TF binding, histone modification, and chromatin structure data could reveal the relative significance of individual features in defining high- and low-expression genes.

We are not the first to build predictive models of gene expression (Table 1), though to our knowledge we are the first to do so in *Plasmodium*. To keep the model simple, we focus on the binary classification task, in which each gene is either “on” or “off,” rather than the more challenging regression setting. We build separate models in three different stages of the *P. falciparum* life cycle and analyze the resulting models to understand which features are implicated in the up- or down-expression of *Plasmodium* genes in different stages of the parasitic life cycle.

**Table 1:**
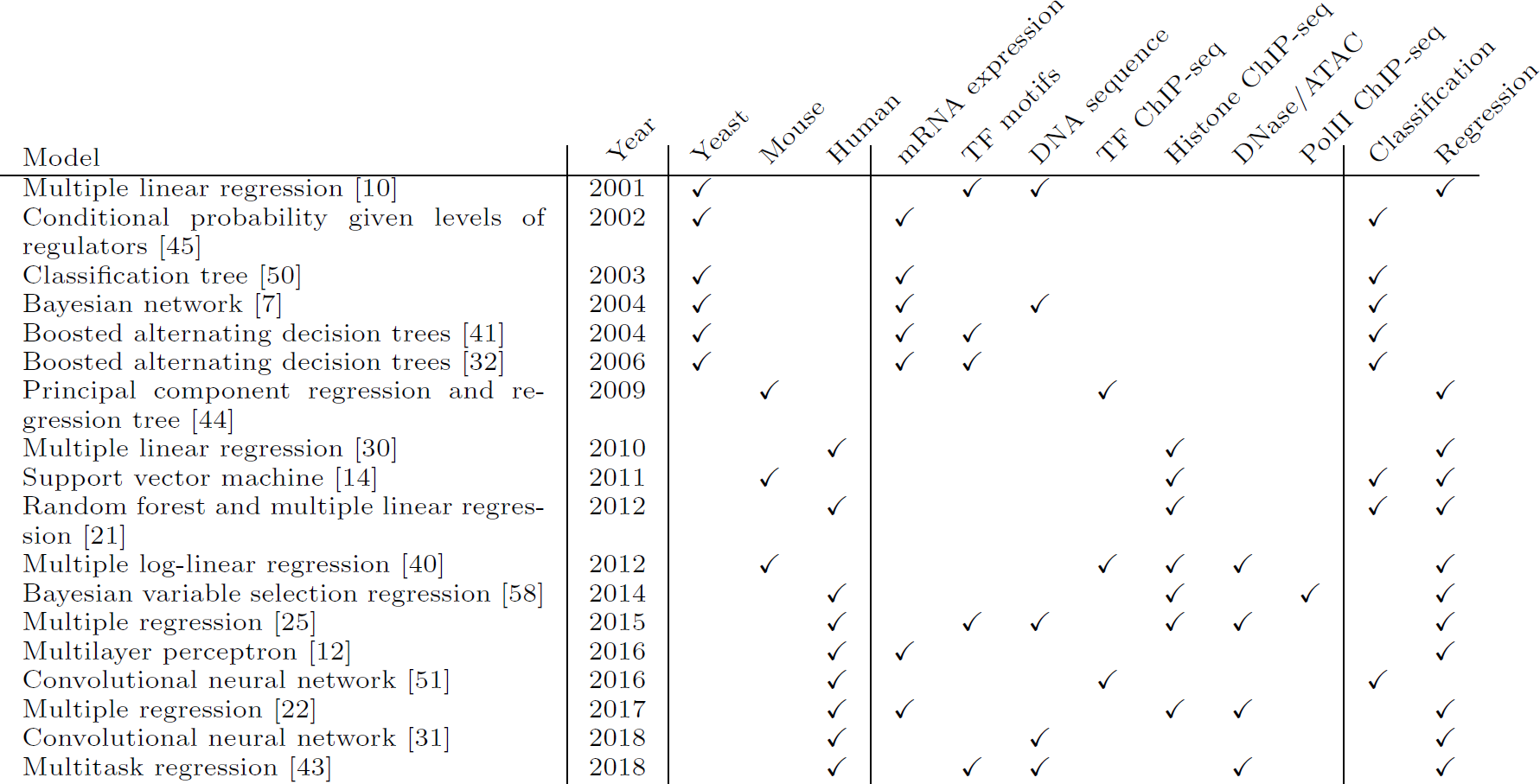
Comparison of methods for predicting gene expression. The “mRNA Expression” feature involves using the mRNA expression of a set of putative regulators to predict the mRNA expression of a different set of target genes.

## 2 Methods

An description of all methods is given below. Code for this manuscript is available online (https://github.com/Daread/plasmodiumExprPrediction).

### 2.1 Data sets

Although *P. falciparum* passes through multiple stages—mosquito, human liver, and human blood—we focus here on the human blood stage of the parasite life cycle, primarily because of the availability of a wide number of relevant data sets. We gathered data for three time points, corresponding to the three main asexual stages within the red blood cell cycle: ring, trophozoite and schizont. Most data sets described below (Table 2) are available in all three of these time points, with the exception of some ChIP-seq data for covalent histone modifications (H3K36me2, H3K36me3, H3K9me3, H4K20me3, and one replicate of H3K4me3 [28]) that were not available for the trophozoite stage.

**Table 2:** Summary of datasets used in classification models. Dataset sources are shown, along with the time points from each study which were used to represent each of three life cycle stages. “N/A” is listed for the GC content and motif score features, as they did not vary across life cycle stages.

| Feature | Study | Description | Ring | Time point |  |
| --- | --- | --- | --- | --- | --- |
|  |  |  |  | Trophozoite | Schizont |
| Distance to telomeres | [2] | Hi-C inferred distance | ✓ | ✓ | ✓ |
| Distance to centromeres | [2] | Hi-C inferred distance | ✓ | ✓ | ✓ |
| Distance to center | [2] | Hi-C inferred distance | ✓ | ✓ | ✓ |
| Nucleosome occupancy | [9] | 100-200 base pair fragments | ✓ | ✓ | ✓ |
| H2A.z | [6] | ChIP-Seq | ✓ | ✓ | ✓ |
| H3K9Ac | [6] | ChIP-Seq | ✓ | ✓ | ✓ |
| H3K4me3 (Bartfai et al) | [6] | ChIP-Seq | ✓ | ✓ | ✓ |
| H3K36me2 | [28] | ChIP-Seq | ✓ |  | ✓ |
| H3K36me3 | [28] | ChIP-Seq | ✓ |  | ✓ |
| H3K9me3 | [28] | ChIP-Seq | ✓ |  | ✓ |
| H4K20me3 | [28] | ChIP-Seq | ✓ |  | ✓ |
| H3K4me3 (Jiang et al) | [28] | ChIP-Seq | ✓ |  | ✓ |
| GC content | See Methods | 100 bp sliding windows | N/A | N/A | N/A |
| TF motifs | [57] | FIMO scan of TF motifs | N/A | N/A | N/A |

#### 2.1.1 Labels and features

##### GRO-seq

To define the on/off labels for our classifier, we used the GRO-seq values from Lu *et al.* [36]. We used GRO-seq values normalized for GC content, gene length, and the “parasitemia factor” of a stage [36], available in Supplementary Table 1. To generate binary labels for genes, for each stage we sorted all protein-coding genes by the normalized GRO-seq counts assigned to that gene in that stage. We labeled the top third of genes as “High expression” and the bottom third of genes as “Low Expression” (Figure 1). The middle third of genes were not used in the analysis.

**Figure 1:**
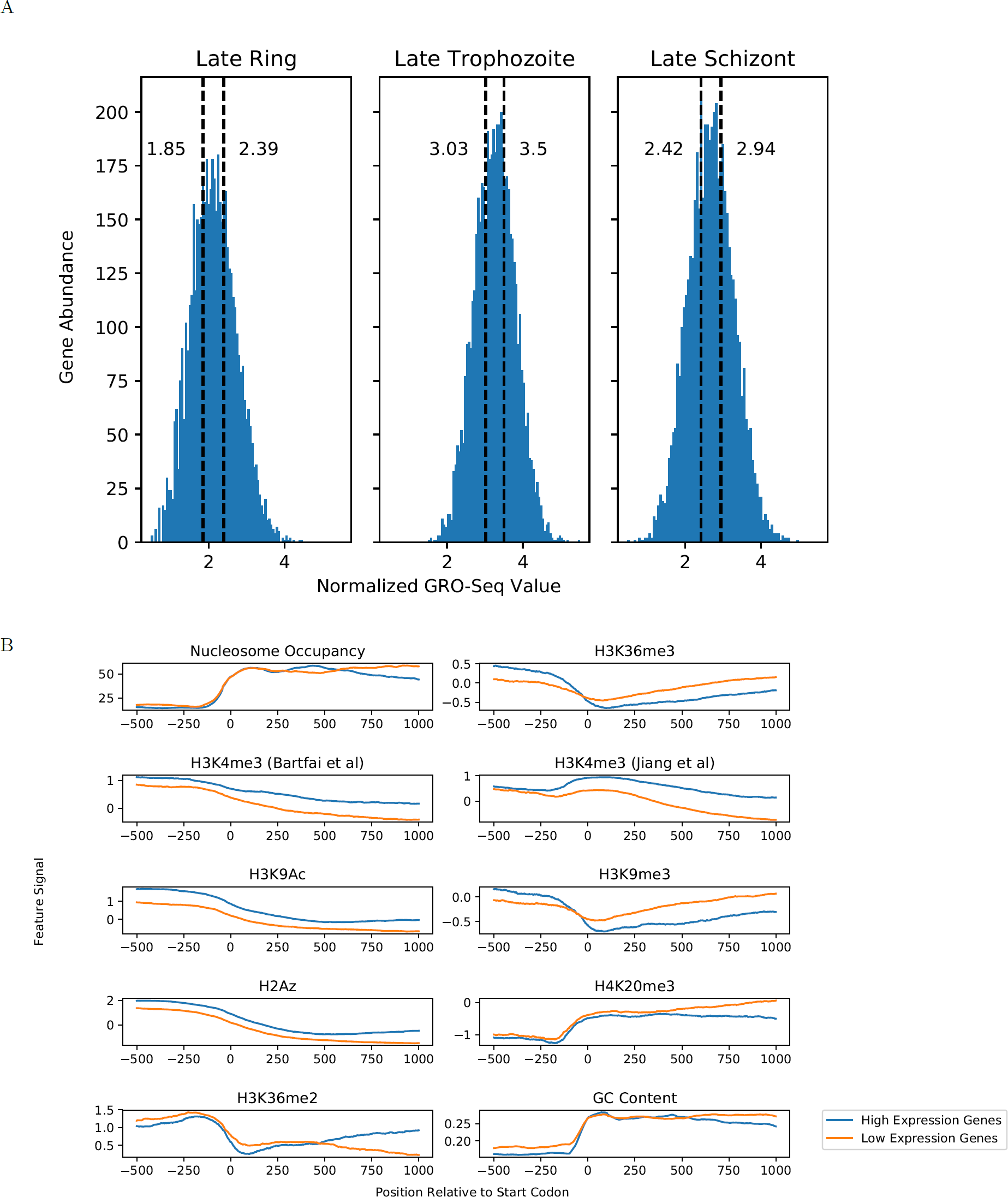
Differences between high- and low-expression genes. (A) Histograms showing the distribution of normalized GRO-Seq values assigned to protein-coding genes within each of three stages of the *P. falciparum* life cycle. The labeled, dashed vertical lines indicate the cut-off values for genes categorized as low- and high-expression. (B) Aggregation plots showing the average signal for features with respect to the start codons of high-expression (blue) and low-expression (orange) genes in the ring stage.

##### Transcription start site and coding sequence annotations

Many of the features that we employed require specifying the start coordinate of each given gene. For this purpose, we use two data sets of co-ordinates: either the start codons from the PlasmoDB v29 annotation or transcription start sites based on CAGE-Seq data from [1]. In that resource, multiple start sites are often annotated for a given protein coding gene. To assign a single TSS for use in feature assignment, we first looked to see if the “primary TSS” assigned in [1] was upstream of the start codon of a gene. If it was, then the start of that TSS was used. Otherwise, we used the TSS lying upstream and closest to the start codon. If no annotated TSS was upstream of the start codon of a gene, then the start codon was used.

##### Histone modification ChIP-seq

ChIP-seq data for the following histone modifications were collected from two studies: H3K4me3, H3K9Ac, and H2.Az from Bartfai et al. [6] and H3K36me2, H3K36me3, H3K9me3, H4K20me3, and H3K4me3 from Jiang et al. [28] Note that one mark, H3K4me3, was measured in both studies. All of the ChIP-seq data was reanalyzed using a standard pipeline that consisted of trimming reads to 76 nucleotides using the fastx toolkit (http://hannonlab.cshl.edu/fastxtoolkit/index.html), mapping reads to the *P. falciparum* genome (PlasmoDB v29) using bwa-mem [33], filtering unmapped and multimapping reads using samtools [34], and generating bedgraph files using bedtools [48]. Fold-change over background was calculated relative to input DNA where available. For histone ChIP-seq datasets from Jiang *et al.*, no input DNA data was available, so values were calculated relative to the mean signal over all the data.

##### Nucleosome occupancy

MNase data from [9] was downloaded in FASTQ format, then trimmed and filtered using sickle version 1.33 (https://github.com/najoshi/sickle). Reads were aligned to the *P. falciparum* genome (PlasmoDB v29) using bwa-mem [33], then sorted and filtered for mapped reads using samtools [34]. A custom Python script selected all alignments between 100 and 200 bp in length, which were then used to generate a bedgraph file using genomeCoverageBed [48].

##### Hi-C

Per-gene distances from telomere centroid, centromere centroid, and center were computed in a previous study [2], using 3D models generated from Hi-C data using PASTIS [55]. These values were obtained directly from https://noble.gs.washington.edu/proj/plasmo3d/.

##### DNA sequence features

Position-frequency matrices for 25 transcription factors for *P. falciparum* were downloaded from CIS-BP [57]. With each motif, we scanned the *P. falciparum* genome using FIMO [26] with a p-value threshold of 0.01. To ensure that the background model represented the unique sequence context of *P. falciparum*, we generated a background model from the *P. falciparum* genome with the MEME Suite command `fasta-get-markov` with Markov order 1 [4]. In addition, percent GC was calculated in 101 base windows centered at each position in the genome.

### 2.2 Predictive model

#### 2.2.1 Models

We used three types of models to classify *Plasmodium* expression and select predictive features. The first was logistic regression with elastic net regularization, using the scikit-learn implementation (sklearn.linear model.SGDClassifier). The second was a tree model with gradient boosting, using the XG-boost Python implementation (xgboost.XGBClassifier). The third was a multi-layer perceptron model, with two hidden layers, each containing the same number of nodes as the input layer. This model was implemented by DeepPINK [37], which is designed to achieve robust feature selection with a controlled error rate.

#### 2.2.2 Performance metric

The performance of each model was evaluated in terms of receiver operator characteristic (ROC) curves. These plots show the rate of true positive classifications (on the y-axis, indicating sensitivity) against the rate of false positive classifications (on the x-axis, indicating 1 - specificity). The area under the ROC curve (AUROC) quantifies the ability of the classifier to balance sensitivity (true positives) against specificity (avoiding false positives). An AUC value of 1 corresponds to perfect performance, whereas a value of 0.5 corresponds to random guessing.

For Figure 2A, the ROC curve for logistic regression classification was generated by combining the gene scores from the test sets in three separate folds of cross-validation. These scores were sorted together to generate the combined ROC curve shown.

**Figure 2:**
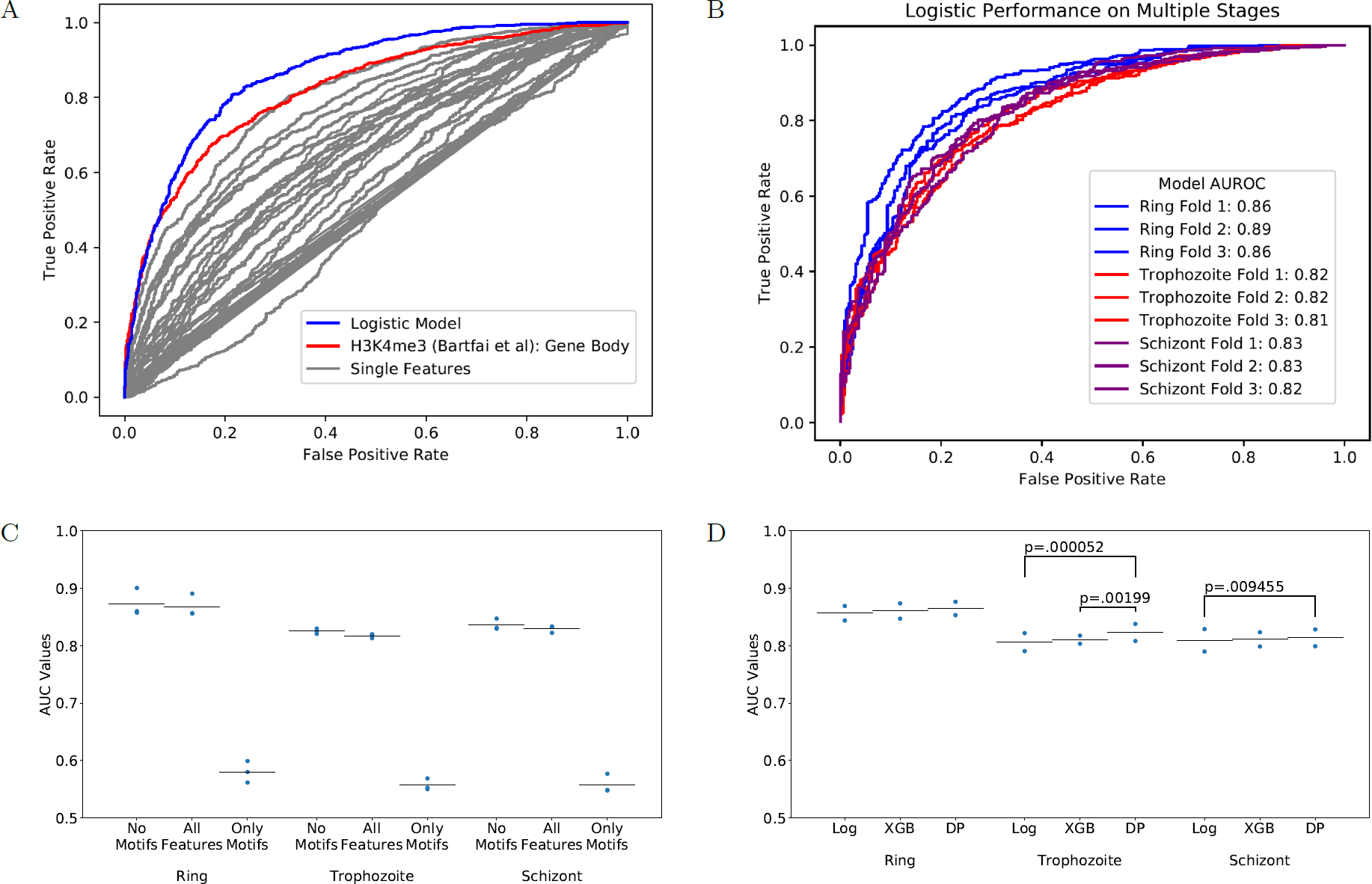
Comparison of classification models. (A) Classification models outperform all individual features for use in classification of gene expression. Grey lines represent the ROC curves resulting from ranking genes by the values of single features in the ring stage, with the best-performing feature shown in red. The blue line represents the ROC curve from training a logistic regression model with elastic net regularization, where the curve is created by combining the predictions across all three test sets. (B) The ROC curves for classification of gene expression by logistic regression across ring, trophozoite, and schizont stages. Individual curves represent performance in one of test three folds in cross-validation. (C) AUROC values for logistic models trained with or without motif scores as features. Points represent AUROC values on the test set in three-fold cross-validation; bars represent average AUROC values on the test data. (D) The AUROC values resulting from training of distinct models in different stages (“Log” = Logistic Regression, “XGB” = XGBoost, “DP” = DeepPINK). Individual points represent the AUROC values from distinct test sets, for the listed model in a given stage. Brackets are labeled with p values for pairwise comparisons within stages where *p* < 0.05, using the DeLong method for comparing AUC values.

#### 2.2.3 Model development

We split the *P. falciparum* into five approximately equally sized (by gene count) folds by chromosome: fold 1 included chromosomes 1, 3, and 13; fold two included 2, 9, and 11; fold three included 7 and 14; fold four included 6, 8, and 10; and fold five included 4, 5, and 12.

The first three folds were used for feature development and hyperparameter tuning. During this stage, we selected hyperparameters by three-fold internal cross-validation. For the logistic regression model with elastic net regularization, we tuned the “alpha” and “l1 ratio” parameters in a sklearn.linear model.SGDClassifier model. For the boosted trees model, we tuned the “max depth”, “min child weight”, “subsample”, “col-sample bylevel”, and “n estimators” hyperparameters in an xgboost.XGBClassifier model. In each case, we performed a grid search across the values listed in Table 3. On the basis of this analysis, we eliminated the motif-based features from our feature set, and we chose to use features based on CDS rather than TSS locations (see Results section for details).

**Table 3:** Hyperparameter selection.

| Model | Parameter | Possible values | Selected value |  |  |
| --- | --- | --- | --- | --- | --- |
|  |  |  | Ring | Trophozoite | Schizont |
| LR | alpha | 0.1, 0.01, 0.001, 0.0001 | 0.0001 | 0.0001 | 0.0001 |
| LR | l1_ratio | 0.4, 0.5, 0.6, 0.7, 0.8, 0.9, 0.95 | 0.95 | 0.8 | 0.9 |
| Boosted trees | max_depth | 4, 5, 6, 7, 8, 9 | 4 | 6 | 4 |
| Boosted trees | min_child_weight | 2, 3, 4, 5, 6, 7 | 5 | 5 | 6 |
| Boosted trees | subsample | 0.3, 0.4, 0.7, 0.8, 1.0 | 0.7 | 0.4 | 0.4 |
| Boosted trees | colsample_bylevel | 0.3, 0.5, 0.7, 1.0 | 0.3 | 0.5 | 0.3 |
| Boosted trees | n_estimators | 40, 60, 80, 100 | 100 | 60 | 80 |

Subsequently, we trained classifiers to make predictions on each of the two test folds, in each case training on the four remaining folds. In this case, hyperparameters were selected that yielded the greatest AUROC value in the three training set “sub-folds” using cross-validation, as implemented in GridSearchCV in sklearn.grid search. The selected hyperparameters are listed in Table 3. Probabilistic classification scores for all genes in both of the two test folds were combined for testing the statistical significance of differences in AUC values. AUC values were compared using the DeLong test for correlated AUCs [20] as implemented in the pROC package in the R language [49].

#### 2.2.4 SHAP Values

The gradient boosting method XGBoost is powerful but challenging to interpret. XGBoost assigns classification labels by taking a consensus decision from an ensemble of individual decision trees [11]. XGBoost models are appealing due to their ability to capture complex interactions among features as well as non-linear relationships between features and classification labels [11]. However, understanding the importance of individual features within such ensembles is challenging, because the model may use a given feature in multiple locations across the individual trees (in contrast to a regression model with a readily interpretable coefficient assigned to a feature).

Consequently, we used SHAP [38] to help interpret the trained XGBoost models. SHAP is a software package that quantifies the effect of each feature on the classification of each example (each gene, in our case) by measuring how much information that feature provides in addition to various subsets of other features being used in the model. Running SHAP on our trained XGBoost models provided us with “SHAP values” for each feature, for every gene. These scores can be studied on a gene-by-gene basis and can be aggregated across all genes within a stage to obtain a consensus score, comparable to a regression coefficient.

#### 2.2.5 DeepPINK

Similar to XGBoost, DeepPINK can also capture non-linear relationships between features and classification labels. Rather than boosted gradients, DeepPINK uses a deep neural network model. Importantly, DeepPINK is able to reliably choose relevant features with a controlled error rate, regardless of arbitrarily complex interactions among features. To rigorously control the false discover rate among selected features, DeepPINK relies upon the recently described model-X knockoffs framework [5]. The primary methodological novelty in DeepPINK is its deep neural network architecture, which enables application of the model-X framework.

### 2.3 Determining feature importance

After training, we examined each model to extract information about which features the model deemed most relevant to the given classification task. For the logistic regression models, we recorded the coefficients assigned to each feature. For XGboost, we used the SHAP package to calculate “SHAP values” for each feature at each gene [38]. The magnitude of the feature importance score was defined as the mean SHAP value across all genes. The sign for the feature importance score (indicating whether a feature indicates high- or low-expression) was defined by the direction of correlation between feature values and SHAP values across all genes. DeepPINK computes feature weights by multiplying the weight matrices across all layers of the deep neural network. The resulting weights can be either positive or negative, indicating the direction of correlation between features and the label. We used the the squared value of the feature weight as the feature importance score.

## 3 Results

### 3.1 Genes labeled as high- and low-expression display differences in several regulatory features

Drawing from a variety of data sources, as described in Methods, we constructed a data set of heterogeneous gene features across each of three stages of the erythrocytic cycle (ring, trophozoite and schizont). GRO-seq measurements of nascent transcription were used to identify genes which high expression (top third) and low expression (bottom third) (Figure 1A).

As an initial step of data exploration, features with signal at a base-by-base level (such as ChIP-seq tracks or GC content) were visualized using aggregation plots showing the average level of signal for a feature with respect to the start codon, segregated by high-expression and low-expression genes (Figure 1B). These plots show expected trends, including enrichment of H3K4me3, H3K9Ac, and H2Az in highly expressed genes, as well as depletion of nucleosome occupancy in promoter regions.

Given the apparent differences in signals upstream and downstream of the start codon, we split each of these main features into two features. The “promoter” feature was the mean feature signal from −500 bases up to the start codon, whereas the “gene body” feature was the mean feature signal from the start codon to 1 kb into the coding sequence. This was done for all covalent histone modification features, H2Az composition, nucleosome occupancy, and GC content.

Similarly, for motif features, we calculated the maximum motif match log-odds score in two windows. The “promoter” window was from −500 bases up to the start codon, while the “gene body” region extended from the start codon up to 1000 bases into the coding sequence.

Ultimately, each ring- and schizont-stage gene was characterized by a set of 73 features, including 50 motif-based features, 14 histone modification features, 7 features characterizing local and global chromatin structure, and 2 features describing local GC content. Trophozoite stage genes used the same feature vector, but with 10 histone features removed due to missing ChIP-seq data sets in that stage.

### 3.2 Machine learning models accurately distinguish between expressed and non-expressed genes

We initially examined single features to establish a baseline of classification performance based on a simple ordering of genes by each individual feature. In this way, we generated one ROC curve for each feature (gray lines in Figure 2A), obtaining AUROCs as high as 0.82 for H3K4me3 gene body signal (blue line in Figure 2A) in classification of ring-stage genes, with similar results obtained in trophozoite and schizont stages (data not shown).

Next, we compared this baseline approach against a machine learning method that integrates all of the available features. We observed, not surprisingly, that an elastic net-regularized logistic regression (“Logistic”) model that integrates all features outperformed rankings based on single features alone: the ROC curve generated by the logistic regression (red curve, Figure 2A) dominates all of the ROC curves generated by ranking genes using single features. We observed similar levels of performance across the three erythrocytic stages, where in a three-fold cross-validated test, the logistic regression model achieved average AUROCs of 0.868 in ring (Figure 2B), 0.817 in trophozoite and 0.829 in schizont.

A key question we aimed to address is the relative utility of the scores derived from TF motifs. Accordingly, we repeated the cross-validated testing of the logistic regression model using three different feature sets: the full set of features, a reduced set in which the TF motif PWM scores have been eliminated, and a set containing only TF motif features. This analysis suggested that the motif features did not aid in classification when combined with non-motif features, and if anything hurt the performance of our models (Figure 2C). Furthermore, models that used only motif features were far less accurate than models that incorporated non-motif features (Figure 2C). Accordingly, motif features were excluded from subsequent analyses.

To determine whether our results thus far depend upon the choice of machine learning method, we also tested two additional types of models: a boosted trees ensemble (“XGBoost”), and a multilayer perceptron with two hidden layers (“DeepPINK”). These methods all demonstrated similar performance on our development folds (data not shown), so we carried all three models into the final testing stage. In this stage, we trained and evaluated each type of model on two folds of data that had not been used in prior model development. The folds each included around 340 high-expression and 340 low-expression genes. In each stage, the three models demonstrate similar AUROC performance, with a slight trend of the multi-layer perceptron model outperforming XGboost, which in turn outperforms logistic regression (Figure 2D). We examined all pairwise model comparisons within each stage (DeLong test for correlated AUCs, see methods), finding three comparisons to have statistically significant differences ((Figure 2D). However, even the differences that are statistically significant are relatively modest in absolute terms, leading us to conclude that each of these machine learning methods achieves reasonably good performance in discriminating between *Plasmodium* genes with high and low expression. Accordingly, we used all three methods in subsequent analyses.

### 3.3 Classification models use stage-specific features

Having established the feasibility of predicting gene expression in *Plasmodium*, we next turned to the more interesting question, namely, which features contribute most strongly to the performance of each classifier? By including three different types of models, we reasoned that if multiple classification models select a similar set of informative features within a single stage, then this would suggest that those features are robust to the choice of model. Accordingly, for each model we calculated a feature importance score (see Methods) on a 0 to 1 scale, where 0 means uninformative and 1 means strongly informative. We also determined the direction of effect, indicating whether a high feature value is predictive of high or low expression. Additionally, the DeepPINK model identifies which features are informative for classification using a method that allows for explicit control of false discovery rate (FDR < 0.05, see Methods). Given the fundamentally different methods used to assign feature importances in each of the three models, we would not expect a precise quantitative agreement in scores (for instance, elastic-net regularized regression models favor zero-valued coefficients, while no such sparisty-inducing behavior occurs in XGBoost or DeepPINK models). Even so, at a qualitative level we clearly observe important patterns in feature importance scores.

Broadly, the three modeling approaches attributed similar levels of importance to features (Figure 3). For example, when assigning importance to histone modifications, the three models tend to agree on which features are informative and their direction of effect: all three models identify high H3K4me3 signal within the gene body as indicative of high expression in the ring stage, select high H3K4me3 signal as indicative of high expression in the trophozoite stage, and identify high H3K9Ac and H4K20me3 signal in the gene body as indicative of high expression in the schizont stage. Similarly, the three models tend to attribute consistent importance to physical chromatin features: all three models highlight the importance of gene body nucleosome occupancy in the trophozoite stage and telomere distance in the ring stage. This consistency across models and methods suggests that our approach to identifying informative features is robust to the differences in modeling approaches.

**Figure 3:**
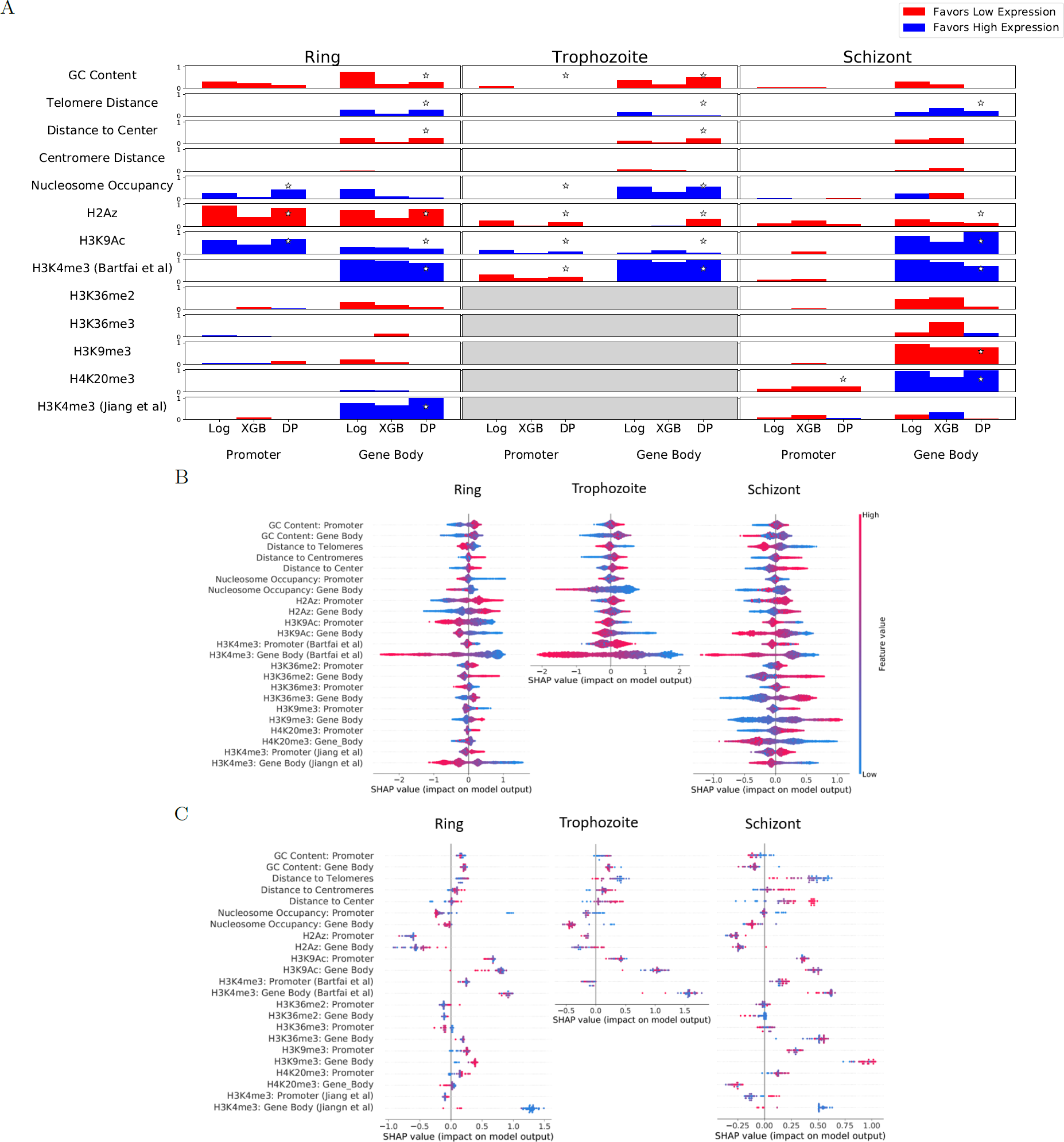
Feature importance measures. (A) Feature importance scores assigned for different models (Log = Logistic Regression, XGB = XGBoost, DP = DeepPINK Multilayer Perceptron). See ”Methods” for details on how feature importance scores were calculated. Scores were normalized to lie within a 0-to-1 range by subtracting the minimum absolute value of all scores in a model, then dividing those numbers by the score with the maximum absolute value. Bar height represents the magnitude of feature significance, while the color of bars indicates the direction of effect (Red: Higher feature value predicts high expression. Blue: Higher feature value predicts low expression.). Features using averages over “promoter” and “gene body” windows (such as ChIP-seq tracks) are split by these sub-features, while features that are not divided (such as distance to centromere centroid) are not. Stars indicate features that were selected as significant using the DeepPINK model, controlling false discovery rate < 0.05. (B) SHAP values for features used in the XGboost classifier for all genes. SHAP values for a given gene represent how significant a specific feature was for classification of a gene as low-expression or high-expression, as well as the direction in which the feature pushed the classification. A positive SHAP score for a feature for a specific gene means that the value of that feature was changed that gene’s classification toward “low expression,” while a negative SHAP value means that feature pushed the gene toward a label of “high expression.” (C) SHAP values for features used in the XGboost classifier for virulence genes.

Interestingly, in the DeepPINK model H2Az coverage in the gene body of trophozoite genes was marked as significant (FDR < 0.05) and associated with low expression. In contrast, scores assigned to this feature were close to zero for both the Logistic and XGBoost models. H2Az signal was previously reported to be almost completely absent from gene coding sequence [6], which makes the apparent significance of gene-body H2Az signal quite surprising. Follow-up validation would be required to see if a minimal level of H2Az truly encodes information within coding sequence, or if the identification of the feature as significant is an artifact of the DeepPINK procedure. However, previous studies in metazoan genomes have also identified H2A.Z in gene bodies. While some research groups link low levels of H2A.Z with inhibition of transcription in reconstituted nucleosomes [39, 53], others suggest that H2A.Z nucleosomes may facilitate transcriptional elongation [56]. Our results support a model in which a low level of H2A.Z nucleosomes acts as a simple barrier to transcriptional elongation. However, given the general agreement between models for almost all other features (Figure 3A) the assignment of importance to H2Az signal by DeepPINK alone suggests that the relationship should be considered very tentative.

In contrast to the high concordance among the three models, we observed low concordance among the importance of individual features across different stages of the erythrocytic cycle. For instance, H4K20me3 was highly informative for predicting a “high expression” label in the schizont stage, but almost completely uninformative in the ring stage (data was unavailable for this mark in the trophozoite stage). The high consistency of feature importance within a stage argues that such differences are not an artifact of the model training process, and suggests that distinct regulatory mechanisms may control transcription in the three different stages. However, further work and replicatation is required to rule out confounding issues such as batch effects between datasets for different stages. For instance, the H3K4me3 features from one source [6] was marked as informative for classification in the schizont stage by all three models, while H3K4me3 signal from a different source [28] was found to be relatively uninformative, by comparison (Figure 3A). Such discrepancies likely stem from differences in either the data generation processes or the synchronization of parasitic stages across distinct sources.

The XGBoost model afforded an additional look at each individual feature’s effects on the classification of single genes. In addition to assigning feature significance and direction-of-effect at the level of the model as a whole (as in Figure 3A), the SHAP score for the XGBoost model can be calculated separately for each feature at each gene. The resulting distribution of per-gene SHAP scores for each feature (Figure 3B) suggests that some features exhibit non-linear relationships between feature score and expression prediction, visually observable as asymmetry in the density plots shown in Figure 3B. For instance, the effect of H3K36me2 gene body signal in the schizont stage model is not a simple relationship where increases in the feature value lead to consistent changes in model classification. This pattern is clearly visible when directly comparing the H3K36me2 gene body signal in schizont genes (x-axis) to the SHAP score of the gene (y-axis) in Figure 4A. Low to intermediate values of H3K36me2 give similar affects upon predictions across a wide range of H3K36me2 signal. However, this trend gives way to a range of higher H3K36me2 values where increases in H3K36me2 are strongly and almost linearly related to changes in classification (as quantified by SHAP values). Such shifts in feature/classification relationships over different ranges of values are observable in several features (Figure 4). This observation suggests that some epigenetic factors influence expression through a relationship more complicated than additive, independent effects.

**Figure 4:**
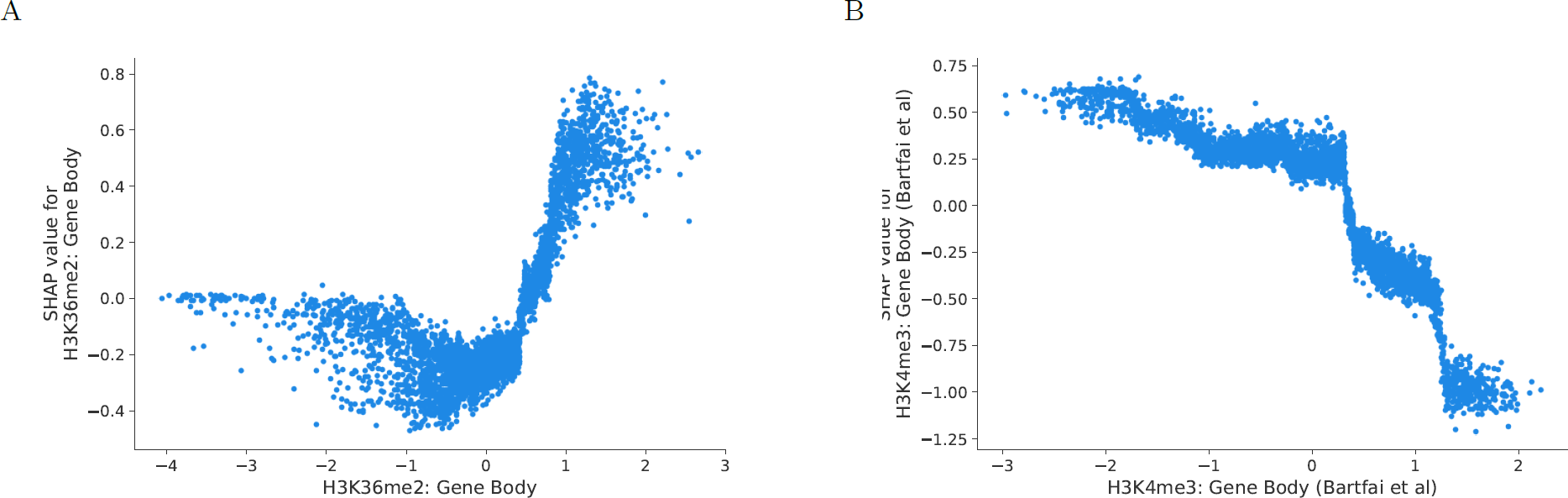
Nonlinear relationships between feature values and SHAP values. (A) Scatter plot showing the values of H3K36me2 signal in the gene body of schizont-stage genes (x-axis) against the SHAP values assigned to those genes (y-axis). (B) A scatter plot plot showing H3K4me3 in the gene body of schizont genes ([6], x-axis) against SHAP values for those genes (y-axis).

We also repeated the per-gene SHAP analysis for the *Plasmodium* virulence genes, which encode a protein family that functions to anchor infected erythrocytes to the endothelium of blood vessels and are an important target for immune recognition [23]. The virulence genes are tightly regulated, with each parasite expressing exactly one of the 60 genes at a given time. In agreement with a known role for H3K9me3 in repression of virulence genes [15], we find that virulence genes have large SHAP scores assigned to H3K9me3 signal. This observation demonstrates that the classification model is not only able to find genome-wide rules for classification, but also selects important features with respect to a specific subset of genes, capturing factors that are known to be important for transcriptional control of that gene family.

### 3.4 Start codons outperform TSSs for predicting transcription

During our exploratory work using the three “development folds” of data, we tested two different approaches for defining the start of a gene: transcription start sites (TSSs) and start codons. Surprisingly, this testing indicated that start codons are more useful than TSSs for defining the division between promoter and gene body for our predictive models. We started by using genome-wide CAGE-seq datasets to define transcription start sites for all genes (see Methods). Plots of feature scores with respect to these two types of “start” positions—start codons (Figure 1B) and TSSs (Figure 5A)—qualitatively showed stronger trends between high- and low-expression genes when defining promoter/gene body splits using start codons rather than TSSs. Furthermore, when we trained classifiers to label genes using features split by either start codon or TSS, the models using promoter/gene body definitions split by start codons consistently out-performed models using TSSs (Figure 5B). Consequently, we focused analyses in this work on models that are split by start codons rather than TSSs.

**Figure 5:**
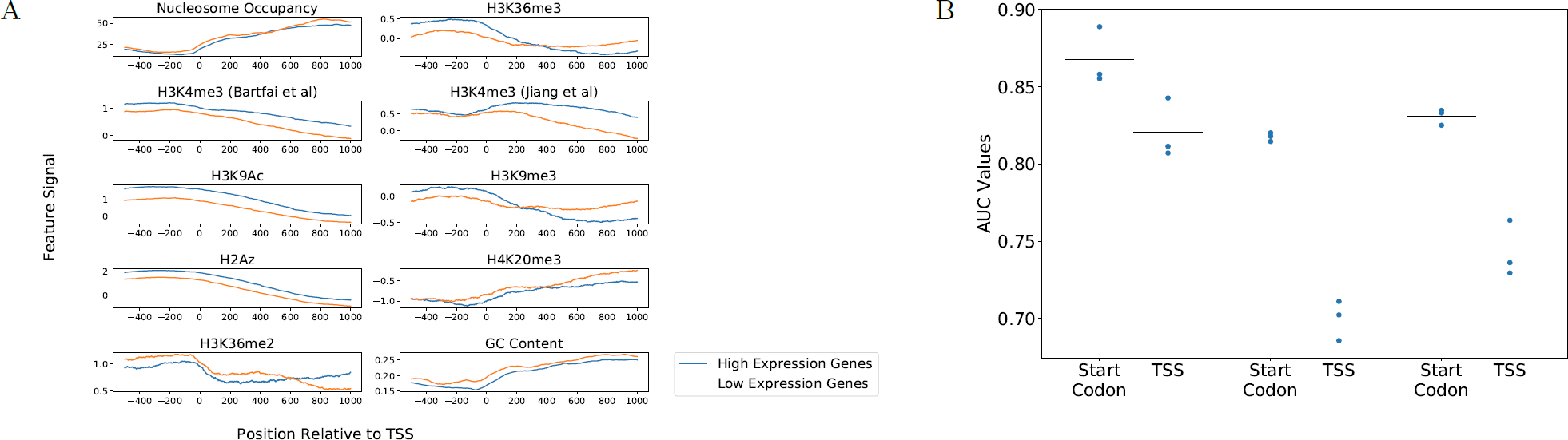
Transcription start sites versus start codons. (A) Aggregation plots showing the average signal for features with respect to the transcription start sites of high-expression (blue) and low-expression (orange) genes. (B) A plot of AUROC scores obtained for training a logistic classifier to classify genes as high- or low-expression. The left column for each stage represents scores using promoter/gene body divided at the start codon (as was done throughout the analysis up to this point), while the right column in each stage used TSSs to divide promoter/gene body regions.

## 4 Discussion

We developed predictive models for *Plasmodium* gene expression that yield AUC values in the range of 0.79– in cross-validated testing. These values are somewhat lower than AUC values reported from studies carried out in other eukaryotes like mouse (0.94 [14]) or human (0.95 [21]). Many factors may contribute to this difference. For example, *Plasmodium* has a smaller number of datasets available for use as features in our models: at most six unique histone covalent modifications were used in our models, whereas 11 unique histone modification features were used in both [14] and [21]. Consistent with this, using a small feature set (five histone modifications) to classify expression in human cells resulted in a model with an average AUC of ∼ 0.8, a value in line with the performance we observed. Furthermore, compared to human and mouse, *Plasmodium* has far fewer genes, which yields fewer examples for training our models. Additionally, the high AT-content of the *Plasmodium* genome presents a consistent challenge to generation of high-confidence genomic datasets [3], so noise in feature datasets may have led to reduced accuracy. An alternate explanation comes from the apparent abundance of genes related to post-transcriptional regulation, rather than genespecific transcriptional control [17]. This discrepancy has led to speculation that the most significant level of gene expression control occurs at regulation of translation, relaxing requirements for strict transcriptional regulatory programs [17]. It is possible that a relatively low reliance on strict transcriptional control allows the parasite to tolerate high noise in transcriptional regulation, in turn leading to a system that is difficult to model accurately.

One surprising outcome was the apparently low utility of features derived from TF binding motifs. This result may be indicative of the relatively low importance of local TF activity in regulating gene expression during the erythrocytic cycle. Alternatively, the result may arise simply because the motifs used here are of low quality or because the way we employed the motifs (by scanning and aggregating p-values) is suboptimal. Previous models of mammalian gene expression based on sequence motifs captured a small amount of gene expression variation [16, 10], while models using TF binding assayed by ChIP-Seq were able to predict expression with far greater accuracy [13]. Clearly, an extensive collection of TF ChIP-seq data would be hugely valuable in exploring the extent to which TFs play an active role in gene regulation in *Plasmodium*. With ChIP-seq data for both histone modifications and TF binding, it would be possible to directly compare the relative importance of specific TF binding versus histone modifications for predicting gene expression.

Inspection of our trained models revealed the use of multiple types of features, from local histone modifications to high-order spatial positioning. Covalent histone modifications were consistently found to be informative features, including the designation of gene-body H3K9Ac and H3K4me3 [6] as statistically significant by DeepPINK FDR control in all three stages (Figure 3A). Furthermore, nucleosome occupancy and GC content were repeatedly identified as informative features (Figure 3A, Ring and Trophozoite feature use). Together, these observations indicate that nucleosome occupancy, histone modification status, and GC content all contain valuable information regarding the activity status of a locus. In addition, the high-order features of gene distances to telomere cluster and nuclear center were also consistently informative for classification of *Plasmodium* gene expression, albeit to a lesser extent than local features such as histone modifications (Figure 3A). This is consistent with previous observations that *Plasmodium* expression correlates with gene spatial positioning [2], and suggests that *Plasmodium* may encode regulatory information in the 3D position of a gene, in addition to its local epigenetic state. Our findings complement previous identification of co-regulatory relationships between functionally related genes in *Plasmodium* [47], with our analysis identifying a repetoire of epigenetic features that underpin such observed patterns.

Some features were consistently informative for classification across the three stages. For instance, high H3K9Ac and H3K4me3 signal in the gene body was indicative of high transcription in every model, for all three stages (Figure 3A). We additionally note that GC content, telomere distance, and distance to center were found to have consistent effects on expression predictions, both between models within a stage and across stages (Figure 3A). This suggests that a number of features included in our models were informative for predicting expression across the intraerythrocytic life cycle.

In contrast to features that displayed consistent effects across stages, some features were only informative in particular stages of the erythrocytic cycle. In this regard, the simplest comparison is between models trained on ring and schizont stage genes, because these stages used the same set of input features. We see a that more histone marks are designated as informative in the schizont stage than in the ring stage, including heavy use of H3K9me3 and H4K20me3 in schizont models (Figure 3A). Meanwhile, the ring-stage model finds H2Az signal and GC content to be more informative than schizont-stage models (Figure 3A). This difference suggests that transcription in the ring stage is related to the incorporation of H2Az histones, GC content, and a small number of histone modifications like H3K4me3. In contrast, in schizont-stage parasites transcriptional activity is linked to a diverse range of covalent histone modifications. Within the models trained to classify trophozoite-stage genes there is a higher use of gene body nucleosome occupancy signal, with respect to ring- or schizont-specific models (Figure 3A), consistent with dramatic changes in nucleosome occupancy during the trophozoite stage being correlated to transcriptional activation [9, 46]. However, the limited data availability for the trophozoite models makes clear interpretation of trophozoite-specific features challenging.

An inherent limitation of our analysis is that, given these data, we cannot easily separate correlations from causative relationships. This is particularly important when modeling transcription using epigenetic data, given previous evidence that some histone marks (H3K36 and H3K79 methylation) are deposited directly as an effect of Pol II elongation, rather than preceding transcriptional activation [24]. Despite this limitation, our identification of predictive features is helpful on two fronts. First, epigenomic changes resulting from transcriptional activity can themselves serve in regulatory roles. In some species, H3K36 methylation, for instance, is deposited concurrently with transcription but serves a regulatory role thereafter, suppressing aberrant initiation of transcription within gene bodies [24]. This means that our models may identify factors important not only for regulation preceding initial activation of a locus, but also for feedback regulatory mechanisms. Second, the observed differences in selected features in distinct stages can potentially prioritize the points in the *Plasmodium* life cycle where experimental dissection of epigentic function would be most informative. For instance, H4K20me3 is not predictive of expression the ring stage, but is consistently associated with transcriptional repression in the schizont stage (Figure 3A). Whether H4K20me3 is a cause or effect of transcription, the molecular events linking this mark to transcription likely only take place— and are experimentally targetable—in the schizont stage. This prioritizes work using schizont-stage cells to interrogate the mechanisms linking transcription and H4K20me3, for instance. Until such follow-up work takes place, however, the predictive relationships that we observe between features and expression cannot be clearly defined as directly regulatory or not.

Analyzing XGBoost models suggested that the best solution to the classification task did not take the simple form in which a unit increase in a given feature leads to a specific, constant change in classification probability (Figure 3B and 4). Consistent with this, our two approaches that allow for feature interactions and non-linear feature/classification relationships, XGBoost and DeepPINK, slightly but consistently out-performed logistic regression (Figure 2D). However, the improvements in test AUC for the XGBoost/DeepPINK models are statistically significant in only a subset of these comparisons, and in all cases are quite modest in absolute terms. This is consistent with work in other eukaryotic genomes, where incorporating feature interactions provided minimal improvement in gene expression prediction accuracy, compared to simple linear models [14, 21]. It is possible that complex models would demonstrate a more significant advantage in a regression task—such as predicting absolute mRNA abundance—rather than the binary classification task that we considered. In our case, however, it appears that models using simple additive effects, such as logistic regression, captured most of the information found within the input features.

During model development and feature refinement, we came to the surprising discovery that placing the promoter/gene body division using start codon position was more effective than using transcription start sites (Figure 5B). This observation is consistent with a previous analysis in which five out of six covalent histone modifications associated with high transcription in *Plasmodium* displayed peak enrichment at the start codons of *Plasmodium* genes, while only one displayed the highest enrichment at transcription start sites [27]. Additionally, this is consistent with the observation that *Plasmodium* lacks a strongly positioned +1 nucleosome at the TSS, but that clearly positioned nucleosomes are observed at the start and end of coding sequences [9, 3]. In the future, it would be interesting to see if epigenetic information related to transcriptional control is truly encoded primarily with respect to start codons, or if technical artifacts due to the extreme AT bias in non-coding DNA upstream of start codons leads to the apparently limited information value of TSS-centered signals.

## Financial disclosure

The funders had no role in study design, data collection and analysis, decision to publish, or preparation of the manuscript.

## Competing interests

The authors have declared that no competing interests exist.

## Acknowledgments

This work was funded by National Institutes of Health awards R01 AI106775 and P41 GM103533.

